# Genetic and ecological drivers of molt in a migratory bird

**DOI:** 10.1101/2022.08.17.504285

**Authors:** Andrea Contina, Christen M. Bossu, Daniel Allen, Michael B. Wunder, Kristen C. Ruegg

## Abstract

The ability of animals to sync the timing and location of molting (the replacement of hair, skin, exoskeletons or feathers) with peaks in resource availability has important implications for their ecology and evolution. In migratory birds, the timing and location of pre-migratory feather molting, a period when feathers are shed and replaced with newer, more aerodynamic feathers, can vary within and between species. While hypotheses to explain the evolution of intraspecific variation in the timing and location of molt have been proposed, little is known about the genetic basis of this trait or the specific environmental drivers that may result in natural selection for distinct molting phenotypes. Here we take advantage of intraspecific variation in the timing and location of molt in the iconic songbird, the Painted Bunting (*Passerina ciris*) to investigate the genetic and ecological drivers of distinct molting phenotypes. Specifically, we use genome-wide genetic sequencing in combination with stable isotope analysis to determine population genetic structure and molting phenotype across thirteen breeding sites. We then use genome-wide association analysis (GWAS) to identify a suite of genes associated with molting and pair this with gene-environment association analysis (GEA) to investigate potential environmental drivers of genetic variation in this trait. Associations between genetic variation in molt-linked genes and the environment are further tested via targeted SNP genotyping in 25 additional breeding populations across the range. Together, our integrative analysis suggests that molting is in part regulated by genes linked to feather development and structure (*GLI2* and *CSPG4*) and that genetic variation in these genes is associated with seasonal variation in precipitation and aridity. Overall, this work provides important insights into the genetic basis and potential selective forces behind phenotypic variation in what is arguably one of the most important fitness-linked traits in a migratory bird.

## INTRODUCTION

Seasonal migration is energetically costly, often requiring extensive morphological and physiological changes each year to prepare for long-distance movements. Molting, defined as the replacement of hair, feathers, skin, and/or exoskeletons to make way for new growth, is one such morphological change that can help prepare animals for long-distance migratory journeys (e.g., smoltification in fish^1^, exoskeleton molting in insects^2^, and feather molting in birds^3^), but can also come at the cost of elevated energetic demands, increased risk of predation, and increased exposure to environmental conditions. In migratory birds, the potential costs associated with premigratory feather molting, a period when feathers are shed and replaced with newer, more aerodynamic feathers^4–6^, are thought to be outweighed by the benefits of increasing flight efficiency during migration^7^. However, feathers are also critical to providing birds with insulation from thermal extremes (either too hot or too cold) and, as a result, avoidance of extreme temperatures during feather molting is important to survival^8,9^. While previous research has demonstrated that intraspecific variation in the timing and location of molt in birds is at least in partly genetically determined^10–12^, very little is known about the genetic basis of feather molting or the environmental factors which select for variation in this key fitness-linked trait.

The timing and location of feather molting in birds is known to vary within and between species in ways that minimize energetic costs and maximize gains of increased flight efficiency. Most migratory passerines complete their molt on the breeding grounds prior to autumn migration, separating the energetically costly stages of migration and molting into different periods of the year. However, some migratory birds have evolved a strategy referred to as molt-migration, where birds first migrate south to take advantage of resource pulses at stop-over locations to complete their molt before heading to wintering areas^13^. One hypothesis proposed to explain the evolution of molt-migratory behavior is the push-pull hypothesis. For migratory birds breeding in Western North America, the push-pull hypothesis posits that molt-migratory behavior evolved in response to migratory birds being pushed away from breeding sites in late summer due to dry conditions and an associated lack of food resources and simultaneously pulled towards monsoon regions of western Mexico to take advantage of pulses in resource availability to complete their molt^14^. Thus, in addition to allowing migrants to take advantage of additional food resources in southern regions, molt-migration may allow birds to avoid exposure to heat stress during the molting period when the temperature regulating benefits of feathers are weakened. If the push-pull hypothesis holds true, we predict that seasonality and changes in precipitation across the breeding range will be positively associated with genetic variation in genes linked to molting, resulting in a higher proportion of molt-migrant associated genotypes in regions characterized by higher seasonality and more extreme summer temperatures.

It has also been hypothesized that intraspecific variation in molting phenotypes (i.e., timing of molt and direction of migration to stopover or wintering grounds) may contribute to the evolution of and the maintenance of migratory divides, regions of overlap between distinct populations with divergent migratory strategies^15,16^, but data supporting this mechanism is limited. For example, differences in migratory direction may result in post-mating reproductive isolation via reduced fitness of hybrids that migrate along intermediate routes^17–20^. Alternatively, hybrids with intermediate molting behavior may suffer reduced fitness if they molt in suboptimal locations^21^. If differences in molting behavior contribute to reproductive isolation, we would expect to see evidence of reduced gene flow associated with transitions between distinct molting behavior, but empirical support for this idea has been limited partially due to difficulties with identifying the molting phenotype of birds captured on their breeding grounds^22^.

While molting strategies have historically been defined on a species level, new research tools for assessing the molt locations of individuals have revealed previously unrecognized levels of intraspecific variation in molting strategies^23,24^. For example, estimates of the geographic area and the environmental characteristics (e.g., type of dominant vegetation, proximity to the ocean, or elevation) of a molting individual can be determined from a single feather using stable isotope analysis (SIA). Once formed, keratinous tissues such as hair, feather or nail are metabolically inert and so their isotopic ratio of common elements such as hydrogen, carbon, and nitrogen, reflect the environmental conditions of where they were grown^25,26^. Thus, SIA has become one of the forefront techniques in avian ecology studies and offers an indirect approach for describing individual variation in the molt strategies^5,27,28^. Now that the research tool is in place for revealing the extent of intra-species variation in molting preferences and environmental conditions, we can begin to ask questions about the genetic and environmental drivers of this complex trait within species.

Next-generation sequencing has facilitated the ability to assess genetic and environmental drivers of complex traits using a variety of techniques that were formerly only available to researchers working on model systems^29,30^. In particular, genome-wide association studies (GWAS) allow for the identification of genetic variants significantly associated with a phenotype of interest^31^, while gene-environment correlation analysis (GEA) analyses are used to identify putative environmental drivers of local adaptation. Given the similar goals of each approach and the fact that natural selection on complex life history traits is generally driven by environmental variation across space, a method that combines the two approaches has the potential to identify environmental drivers of a specific complex phenotype rather than environmental drivers of local adaptation more generally. Here we adopt a two-step approach that first uses GWAS to identify candidate loci underlying molt-migratory behavior in a migratory songbird, the Painted Bunting (*Passerina ciris*), and then uses GEA to identify environmental drivers of genetic variation linked to this phenotype.

The Painted Bunting is a songbird which breeds in two disjunct populations across southwestern United States (U.S.) and northwestern Mexico (larger western population) and along the Atlantic coast of the U.S. (smaller eastern population) from Florida to North Carolina^32^. It is an ideal system in which to investigate the genetic and ecological drivers of intraspecific variation of molt because field observations, tracking studies, and isotope analysis have documented clear differences in molting strategies across the range^33–35^. More specifically, previous work has shown that southwestern breeding birds are molt-migrants that stopover in western Mexico in the fall to complete their molt, while eastern populations follow the more common strategy of molting at the breeding grounds before migrating to southern latitudes for the winter^33,36,37^. However, within the southwestern region, the variation of the molt-migratory phenotype and its potential role in local adaptation and gene flow between populations exhibiting distinct molting phenotypes have not yet been described. Here we identify variation in hydrogen stable isotope values extracted from feathers as a proxy for environmental molting conditions experienced by individuals sampled across the breeding range (e.g., environments near the breeding ground or farther away) and then pair this approach with GWAS to reveal genetic variation associated with molting behavior. We then integrate GWAS and GEA analyses to test whether the environmental drivers of molt-linked genetic variation are in keeping with the pushpull hypothesis.

## METHODS

### Sample collection and DNA isolation

We compiled DNA samples from 192 individuals from 13 populations across the Painted Bunting’s breeding range. At each site, birds were captured using targeted mist-netting and blood samples were collected via brachial venipuncture and preserved in Queen’s lysis buffer^38^. Importantly, population sampling for genetic analysis focused on 13 populations spanning a range of hypothesized differences in molting behavior of the Painted Bunting (Figure 1; Table S1). Further, we included additional 261 genetic samples from 13 sites that overlap with the RAD-seq data set in addition to 12 new breeding populations to validate associations between allele frequencies and environmental variables in key candidate loci identified via RAD-seq alone. DNA from all samples was purified using the Qiagen™ Dneasy Blood and Tissue extraction kit and quantified using the Qubit® dsDNA HS Assay kit (Thermo Fisher Scientific). Remaining blood samples were cataloged and stored for future use in -80°C freezers at the Conservation Genomic Laboratory at Colorado State University.

**Figure 1.**
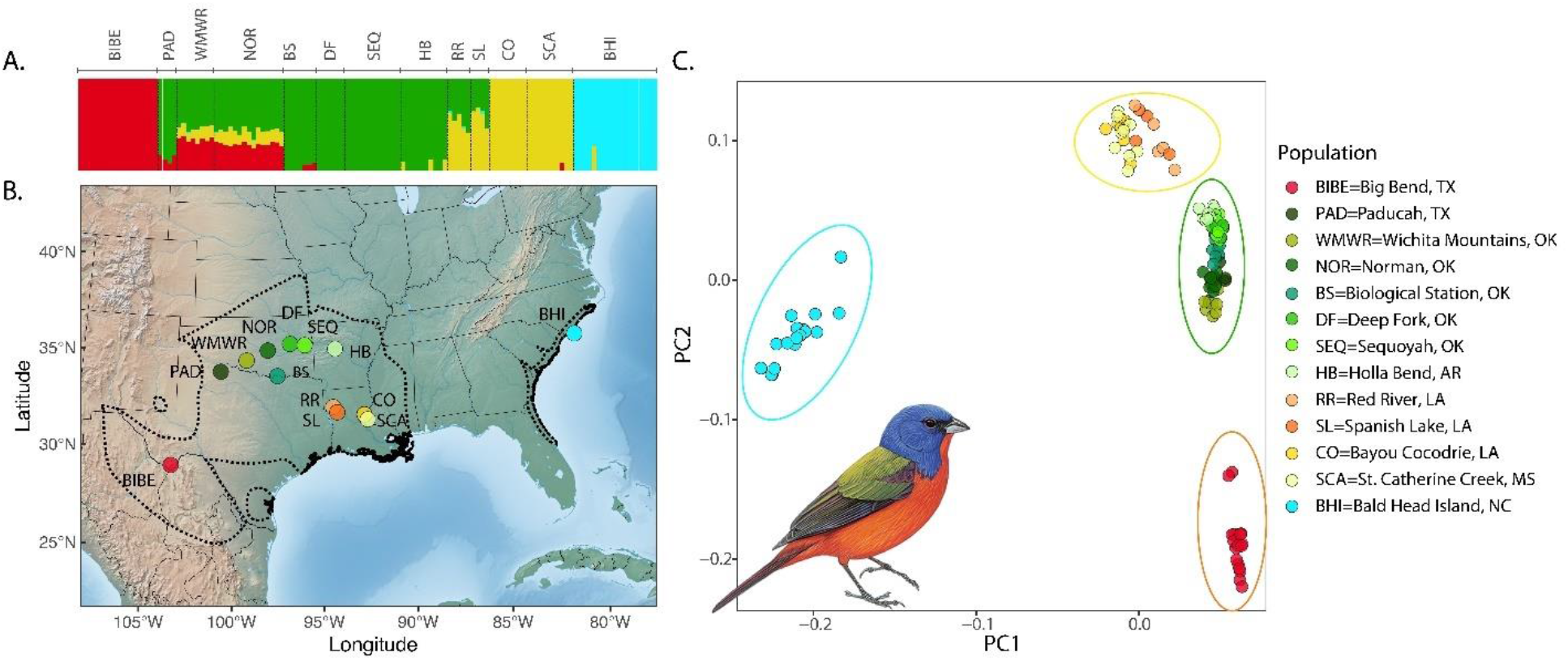
The breeding range of Painted Buntings used for this study, and the genetic clustering of sampled individuals. A) The sampled individuals represent four distinct genetic clusters (Texas: red, Central: green, Louisiana: orange/yellow, and Coastal clusters: cyan). B) Breeding individuals were sampled from 13 locations spanning the entire breeding range of Painted Buntings (dotted line). C) PCA of genetic variation of 41,786 variants. Green dots correspond to the Central genetic cluster and our sampling transect across populations with differences in molting behavior.

### RAD sequencing

Genome scans were conducted using high-density RAD-Seq on all 192 individuals following a modified version of the bestRAD library preparation protocol^39,40^. In short, DNA was normalized to a final concentration of 100ng in a 10ul volume, digested with restriction enzyme SBfl (New England Biolabs, NEB). The fragmented DNA was then ligated with SBfI specific adapters prepared with biotinylated ends and samples were pooled and cleaned using 1X Agencourt® AMPure XP beads (Beckman Coulter). Pooled and clean libraries were sheared to an average length of 400bp with 10 cycles on the Bioruptor NGS sonicator (Diagenode) to ensure appropriate length for sequencing and an Illumina NEBNext Ultra DNA Library Prep Kit (NEB) was used to repair blunt ends and ligate on NEBNext Adaptors to the resulting DNA fragments. Agencourt® AMPure XP beads (Beckman Coulter) were then used to select DNA fragments with an average length of 500bp, libraries were enriched with PCR, and cleaned again with Agencourt® AMPure XP beads. The resulting libraries were sequenced on two lanes of an Illumina HiSeq 2500 at the UC Davis Genome Center using 250 bp paired-end sequencing.

We used the program *stacks*^41^ to demultiplex, filter and trim adapters from the data with the *process_radtags* function and remove duplicate read pairs using the *clone_filter* function. We mapped the processed sequences to the annotated genome of a closely related relative, the Medium Ground finch (MEGR), *Geospiza fortis*^42^ (NCBI Assembly ID: 402638 (GeoFor_1.0)). This genome is from a female individual sequenced at 115X coverage with HiSeq data. The genome is 1.07 Gb, and scaffold N50 is 5.2 Mb. We mapped the processed reads to the MEGR genome using *bowtie2*^43^ and detected variants using the program HaplotypeCaller in the Genome Analysis Toolkit (gatk)^44,45^. For initial filtering, we used *vcftools*^46^ to remove indels, non-biallelic SNPs, and low quality and rare variants (genotype quality 20; coverage depth 10; minor allele frequency 0.03). This initial filtering resulted in 86,347 loci in 192 individuals. The final number of SNPs and individuals to be retained for the subsequent analyses was assessed by visualizing the tradeoff between discarding low-coverage SNPs and discarding individuals with missing genotypes using custom scripts within the R-package *genoscapeRtools*^47^. The final dataset included 41,786 variants in 124 individuals.

### Population structure

To determine whether population structure was associated with a transition between molting phenotypes, we assessed patterns of population structure using the program ADMIXTURE^48^, for K = 1 to K = 6 putative clusters, with a model that accounted for admixture between populations and correlated allele frequencies. We ran 5 iterations for each value of K, with a burn-in period of 50,000, and a total run length of 150,000 generations. To determine the optimal number of genetic clusters we used the cross-validation method, a process of systematically withholding data points to identify the best K value^49^. We used this algorithm to detect the uppermost hierarchical level of structure across the Painted Bunting breeding range and visually inspected subsequent structure plots using *pophelper*^50^ in R to identify regions where geographic barriers to gene flow exist and/or where admixture homogenizes population structure.

### Stable isotope analysis

To identify clusters of individuals based on their stable isotope values of hydrogen (δ²H) in feathers during molting, we collected wing feathers from 166 birds of which 114 individuals were also sequenced. We washed the ninth primary wing feathers in a 2:1 solution of Chloroform-Methanol following the protocol detailed in Chew et al. (2019). This step and additional washes in a 30:1 solution of deionized water and detergent ensured that debris and oil contaminants were removed from the samples. Feather washing was followed by a drying step of 48 hours at room temperature and cutting to about 200 µg (±10 µg) of the distal feather tip packaged into a silver capsule. Before mass spectrometry analysis, we stored our feather samples for a minimum of three weeks to allow for equilibration of exchangeable H to the laboratory environment (Wassenaar and Hobson 2003).

We ran all the samples through a Thermo Scientific™ TC/EA high temperature conversion elemental analyzer interfaced with a Thermo Scientific™ Delta V Advantage Isotope Ratio Mass Spectrometer via a Thermo Scientific™ ConFlo IV Continuous Flow Interface. We report δ²H values as mean ± SD in delta notation of parts per million (‰) from the standards (δ²H_sample_ = [(*R*_sample_/*R*_standard_) − 1]) comparative to the Vienna Standard Mean Ocean Water [VSMOW]. We used Caribou Hoof Standard (δ²H_CHS_ = -197.0 ‰) and Kudu Horn Standard (δ²H_KHS_ = -54.1 ‰) as external calibration standards (USGS - United States Geological Survey, Reston Stable Isotope Laboratory, Virginia, USA) and brown-headed cowbird feathers (δ2H_BHCO_ = -40.9 ‰) as an internal blind standard. These standards were run after every 10 samples, and all samples were analyzed across three separate runs on the instrument. The absolute errors (‰) for all three standards across all runs (n = 12 combined) were: CHS, mean = 1.3, SD =1.3; KHS, mean = 1.3, SD = 0.6; and BHCO, mean = 1.4, SD = 1.2.

To explore variation in molting strategy across individuals, we extracted stable isotope values of hydrogen in precipitation (δ²H_p_) at each sampling site^51^ and then subtracted the stable isotope values of hydrogen in feathers (δ²H_f_) collected from each individual at that site. Thus, we computed a simple differential index (δ²H_diff_) to estimate whether the keratinous tissue of each individual feather was grown at or nearby the sampling location:

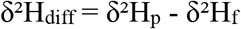

We assumed that δ²H_diff_ values close to zero indicated molting near the sampling location, while δ²H_diff_ values diverging from zero indicated molting farther away from the sampling site.

### Genome-wide association analysis (GWAS) of molting phenotype

We used two genome-wide association study (GWAS) analyses to identify loci associated with molting phenotypes defined by stable isotope groupings. We used a 13.7‰ cutoff to delimit six groups ranging from individuals with extremely negative δ²H values to less negative or nearly positive δ²H values, corresponding to birds molting in different environments. The resultant clusters derive from an ad hoc procedure, but they are centered on differences of more than 12‰ which has been recognized to be ecologically meaningful^25,34,52^. After the samples were grouped based on isotopic differences, we conducted a Bayesian sparse linear mixed model in GEMMA (*bslmm*^53^) to identify single-gene effects. Given we do not know the genetic basis of molting phenotypes, the adaptive *bslmm* model allows us to infer it from the data. *Bslmm* combines both the linear mixed models (*lmm*^31^) and Bayesian variable regression models (*bsvr*^54^), giving the benefits of each when the underlying genetic basis of the trait (*i*.*e*. many genes of small effect vs. few genes of large effect) is unknown^53^. To statistically control for population structure in GEMMA^55^, we incorporated a genetic kinship matrix generated using 41,734 analyzed variants, where missing genotypes were imputed using *beagle*^56^. We then ran the *bslmm* model with the kinship matrix, sampling for 5 million generations and a burn-in period of 500,000 iterations.

Candidate molt-migration associated SNPs were retained when they had a posterior inclusion probability (PIP) threshold > 0.1 ^57^. Second, we employed a multi-locus GWAS method, FASTmrMLM in the program mrMLM^58^, which is designed to detect multiple loci while reducing the chance of false positives. We implemented FASTmrMLM in the program mrMLM using the kinship matrix to account for population structure, and distance between loci to account for interaction among loci given a certain distance (default = 20kb, reported 100kb). To calculate distance between loci, we placed the scaffold positions in the Zebra Finch (*Taeniopygia guttata*) chromosomal order using satsuma2 synteny^59^ prior to the analyses. Candidate molt-migration associated SNPs were those that were found to have a LOD score > 3. To identify genes located near outlier SNPs generated using both single-locus and multi-locus methods, we downloaded Ensembl gene predictions for medium ground finch (GeoFor_1.0, annotation version 102). We used *bedtools* -closest to find the gene closest to the candidate variants detected^60^. We then retained only genes which occurred within a 50kb of the candidate loci.

### Gene-environment correlation analysis (GEA)

To identify the environmental variables that best explained genetic variation underlying the molting phenotype we used gradient forest analysis with the R package *gradientForest*^61^. Climate and environmental data consisted of 19 WorldClim^62^ variables, as well as vegetation indices (NDVI and NDVIstd, Carroll et al. 2004; Tree Cover, Sexton et al. 2013), elevation data (http://www.landcover.org), and a measure of surface moisture characteristics (QuickSCAT from http://scp.byu.edu). The genetic data consisted of allele frequencies from the 412 candidate loci with non-zero effects identified via our single-locus GWAS analysis. To provide a ranked environmental variable list based on the relative predictive power of all environmental variables, we ran *gradientForest* over 100 trees without binning the data due to the small number of candidate loci. To visualize the predicted associations between genetic variation between genetic variation in molt-linked loci and environmental variation across space we used the resulting gradient forest model to predict the association between environmental variables at 100,000 random points across the Painted Bunting breeding range and plotted the relationships using principal components analysis (PCA). To visualize the different adaptive environments across the breeding range, we assigned colors to the breeding range based on the top three principal components axes^61^. To determine whether our results were significantly different from random, we compared our observed results with those obtained from gradient forests run by randomizing the match between allele frequencies and the environmental data (n = 1000).

To further test the relationship between allele frequencies at the top ranked candidate genes identified in the GWAS above and the three highest ranking environmental variables identified by the gradient forest analysis, we used Pearson correlation (FDR corrected p-value <0.05).

### Validating gene-environment relationships in key candidate loci

To validate the association between allele frequencies in molt-linked loci and environmental variation, we created custom assays to genotype 261 new genetic samples from 25 breeding populations (12 new sites and 13 that overlapped with the RAD-seq analysis) at the top 6 candidate genes with the highest association. We used the R package *snps2assays*^63^ to create designable primers (e.g., GC content was less than 0.65, no insertions or deletions (indels) within 30bp, and no additional variants within 20bp of the targeted variable site). Additionally, we aligned 25 bp surrounding the targeted variable sites to the genome using *bwa*^64^ to confirm the primers mapped uniquely to the reference genome. Only the assay targeting *GLI2*, the top ranked candidate loci, passed all filters, and was subsequently genotyped on a Fluidigm™ 96.96 IFC controller. Allele frequencies were calculated for each location after removing individuals with greater than 20% missing data. Standard linear regression was used to test for associations between environmental variation at the top three climate variables and allele frequencies at *GLI2* in both the original (RAD-seq) and validation (SNP-only) datasets (FDR corrected p-value <0.05).

## RESULTS

### Population genetic structure

Overall, analysis of 41,786 total variants identified via RAD-sequencing revealed significant levels of population structure across the Painted bunting breeding range (Fig 1A-B). The PCA plot shows four distinct genetic clusters: an Eastern cluster (North Carolina), a Southwest cluster (Big Bend, TX), a Central-Southeastern cluster (Louisiana), and a Central cluster (including populations from western Oklahoma to Arkansas; Figure 1C). PC1 explains most of the variance (6.36%), separating the eastern genetic cluster from the larger continuous breeding range in the west, while PC2, which explains 2.22% of the variance and separates the southwest cluster (red; Big Bend, TX) from the central and south-central clusters. The admixture analyses support weak population structure between central, southwestern, central-southeastern, and eastern populations, but this may in part result from a lack of sampling at intermediate sites, particularly between sites in the Southwestern and Central regions. While overall genetic breaks within the breeding individuals were not concordant with hypothesized transitions in molting behavior, the population structure results can be incorporated into the subsequent GWAS analyses to reduce false positives related to population-specific genetic differences.

### Stable isotope assignment of molting phenotype

The analysis of hydrogen stable isotope values from feathers (δ²H_f_) showed high variation ranging from -92.9‰ to -10.8‰. The differential index of hydrogen stable isotope values (δ²H_diff_) revealed limited variation at the extreme western (e.g., southwestern Texas) and eastern (e.g., Louisiana, Mississippi, and North Carolina) sampling sites of the breeding range and a larger gradient of values moving from the interior western population (e.g., Oklahoma) towards southeastern U.S. (Fig 2 panel A–C; Table 1). Even though the resolution of our stable isotope analysis is coarse, it is nevertheless indicative of a transition around a -93°.5’ longitudinal meridian, from molt-migrants in the central portion of the western population to individuals that molt at or near their breeding grounds in the east, which is in agreement with previous work based on field observations and tracking data^33^.

**Figure 2.**
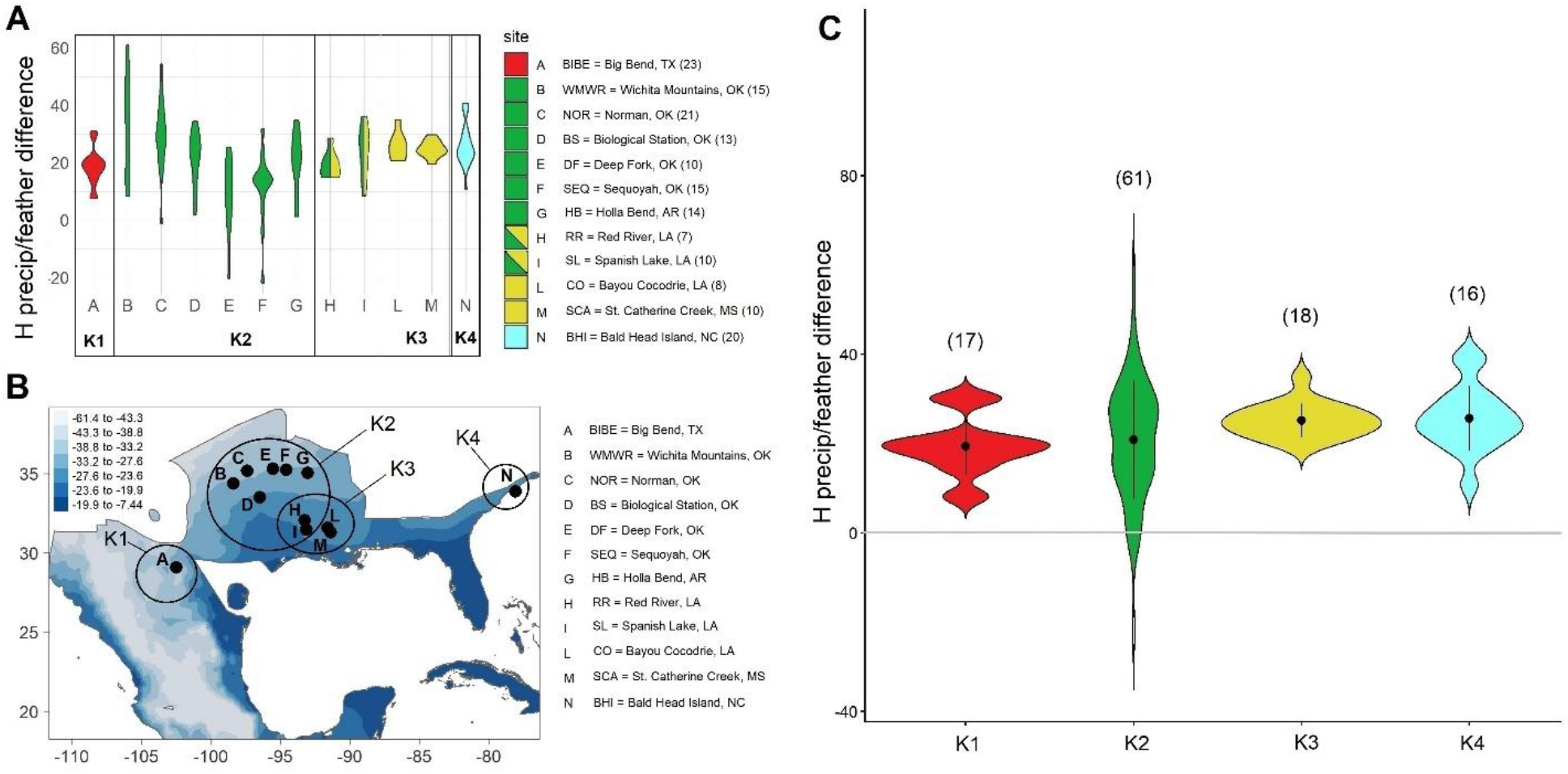
The variation of hydrogen stable isotope values across the breeding range of the Painted Bunting. Different colors match the clusters of the genetic analysis presented in Fig 1. These genetic cluster (K1-4) correspond to an Eastern cluster (North Carolina; cyan), a Southwest cluster (Texas, red), a Central-Southeastern cluster (Louisiana, yellow), and a Central cluster, including populations from western Oklahoma to Arkansas (green). Sample sizes are presented in parentheses. A) Violin plots resulting by subtracting stable isotope values of hydrogen in feathers (δ²H_f_) from stable isotope values of hydrogen in precipitation (δ²H_p_) at each sampling site. δ²H_diff_ values close to zero indicate molting near the sampling location. The largest δ²H_diff_ variation is found in the populations of the Central cluster (green). Samples from sites H and I could not be genetically assigned with certainty and are represented with two colors (green and yellow), to indicate that could be grouped to either cluster K2 or cluster K3. B) Map of hydrogen stable isotope values in precipitation (δ²H_p_) across the Painted Bunting range and sampling locations. C) Violin plots of δ²H_diff_ values of birds grouped in four distinct clusters based on genetic membership (K1-4) and for which individual δ²H_f_ values were also available. As shown in panel A, the largest δ²H_diff_ variation occurs in the Central cluster populations (green). A detailed list of samples used to generate the plots in panel A and C is included in the supplementary material (Table S2 and Table S3).

**Table 1.** Summary table of δ²H_diff_ range variation across sites, including sample size (n), mean and standard deviation. At each site, the δ²H_diff_ index was calculated by subtracting stable isotope values of hydrogen in feathers (δ²H_f_) from stable isotope values of hydrogen in precipitation (δ²H_p_). Smaller δ²H_diff_ ranges (close to zero) indicate molting near the sampling location. The largest δ²H_diff_ variation is found in the populations of the Central cluster (the last six sites at the bottom of the table from BS to NOR).

| site name | State | site ID | n | mean | sd | range |
| --- | --- | --- | --- | --- | --- | --- |
| St. Catherine Creek | Mississippi | SCA | 10 | 24.72 | 2.9 | 10.2 |
| Red River | Louisiana | RR | 7 | 19.49 | 4.75 | 13.6 |
| Bayou Cocodrie | Louisiana | CO | 8 | 25.75 | 4.71 | 14.3 |
| Big Bend | Texas | BIBE | 23 | 18.52 | 6.17 | 23.3 |
| Spanish Lake | Louisiana | SL | 10 | 24.03 | 8.71 | 27.5 |
| Bald Head Island | North Carolina | NC | 20 | 25.92 | 7.38 | 29.8 |
| Biological Station | Oklahoma | BS | 13 | 21.79 | 9.27 | 32.4 |
| Holla Bend | Arkansas | HB | 14 | 20.43 | 10.09 | 33.5 |
| Deep Fork | Oklahoma | DF | 10 | 9.24 | 13.49 | 45.5 |
| Wichita Mountain | Oklahoma | WMWR | 15 | 32.97 | 17.41 | 52.7 |
| Sequoyah | Oklahoma | SEQ | 15 | 11.55 | 12.2 | 53.5 |
| Norman | Oklahoma | NOR | 21 | 29.89 | 11.62 | 55.4 |

### Genes associated with the molting phenotype

Both single gene (*bslmm*) and multi-locus (*mrMLM*) GWAS algorithms were used to find regions of the genome associated with molting, while accounting for population structure. In the *bslmm* analysis, a median of 91.1% of phenotypic variation was explained by the genotype (95% CI 45.2–99.9%), of which 46.7% was explained by SNPs with nonzero effects, but the credible intervals on this estimate were very high (95% CI 2.3–94.4%). Approximately 0.16% of the variants had non-zero effects (n = 412), however 67 were considered to have major effects. We defined the top candidate SNPs as those that were found with sparse effects in at least 10% of the MCMC runs (PIP > 0.1), after controlling for population structure (Fig. 3A). Of these outlier SNPs, the top two were also identified in the mrMLM analysis (Table 2 and Table 3). The strongest single-gene effect was found for a SNP on chromosome 7, within the coding region of the *GLI2* gene. The next two strongest associations were identified on chromosome 4A, situated within the coding region of an uncharacterized protein (LOC102037655), and on chromosome 2, located in the regulatory region approximately 15 kbp downstream from the *HEPACAM2* gene. The multi-locus approach identified 5 additional genes associated with the molt-migration strategy (Fig 3). Similar to the *bslmm* analysis, the highest effect identified in the multi-locus analysis points to a variant in the *GLI2* gene. The remaining genes identified with high effect sizes were *CSPG4, PDGF-B, CNOT9, ARHGAP26*, and two uncharacterized proteins (LOC102037655 and LOC102037327).

**Figure 3.**
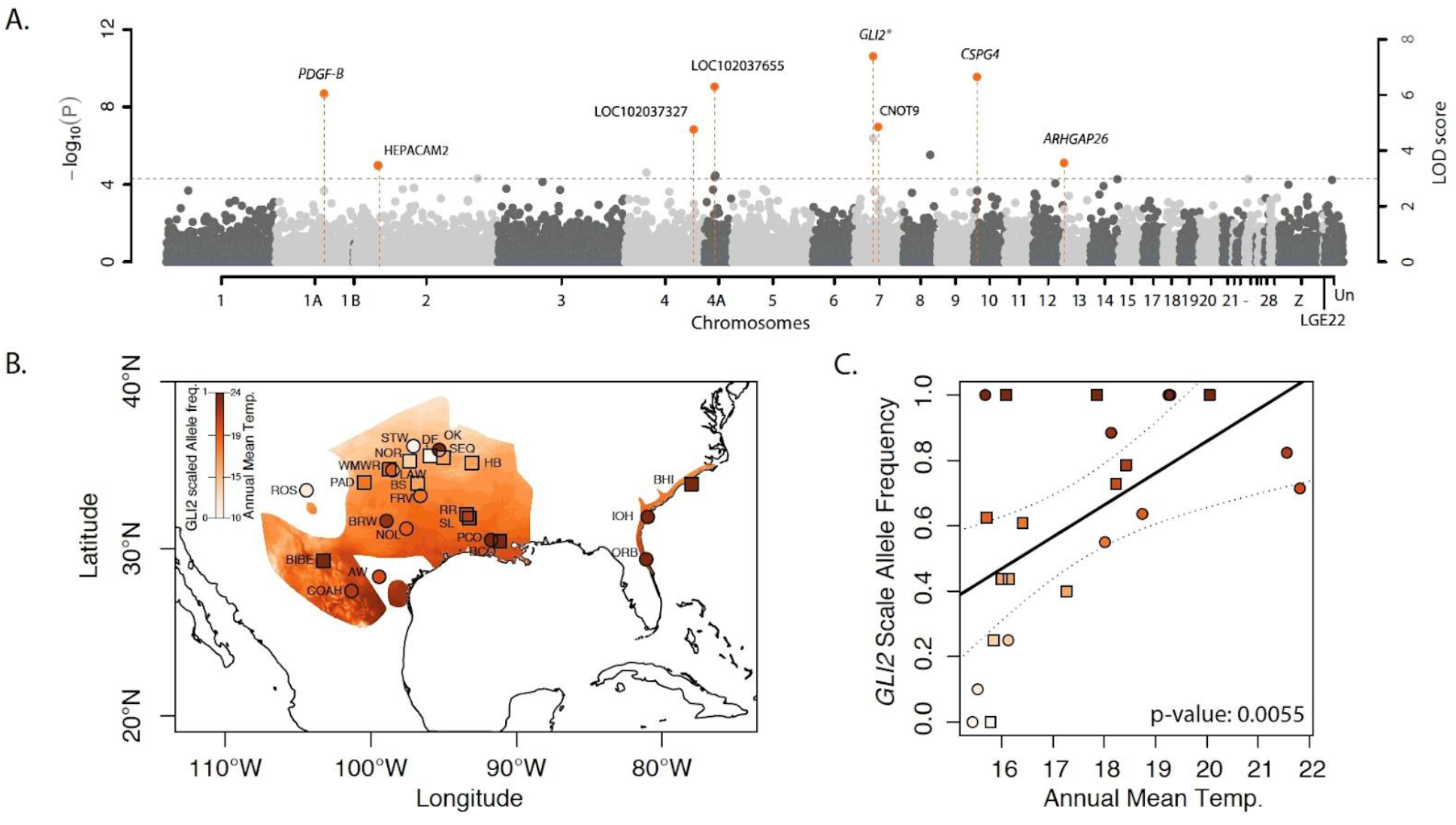
A) Manhattan plot of genome-wide association results of molt-migration behavior grouping in Painted Buntings using the mrMLM v4.0 software. Orange highlights the candidate genes around quantitative trait nucleotides (QTNs). Alternating grey and black colors correspond to alternating chromosomes. (B and C) Correlations between allele frequency and BIO01 (Annual Mean Temperature) for top ranking SNP in *GLI2*. Samples genotyped by RAD-Seq are represented as squares, and samples genotyped with Fluidigm assays are shown as circles. Site abbreviations used on the map in panel B and additional sample information are reported in the supplementary material, Table S4.

**Table 2.** Candidate genes identified in two GWAS analyses. Chromosome and position, gene name and region and distance to region, and parameters estimated using the two GWAS methods (single gene effects: *bslmm* and multigene effects: FASTmrMLM), with their effect sizes, and functional annotation.

| Chromosome | Marker position (bp) | Gene name | Region | Distance to Region (bp) | GEMMA Effect size | FASTmrMLM |  |  |  | Function |
| --- | --- | --- | --- | --- | --- | --- | --- | --- | --- | --- |
|  |  |  |  |  |  | QTN effect | LOD score | -log10(P) | r<sup>2</sup> (%) |  |
| 7 | 15411182 | GLI2 | intron 12 | 0 | 0.12 | 0.87 | 7.40 | 8.27 | 24.82 | Functions as transcription regulator in the hedgehog (Hh) pathway |
| 10 | 3306352 | CSPG4 | exon 7 | 0 | na | -0.35 | 6.65 | 7.51 | 11.55 | Stimulates endothelial cells motility during microvascular morphogenesis. Cell surface receptor for collagen alpha 2(VI) |
| 4A | 8965887 | LOC102037655 | intron 2 | 0 | 0.09 | -0.29 | 6.30 | 7.15 | 7.65 | Protein FAM162B-like |
| 1A | 50451000 | PDGF-B | Regulatory 1 | 2502 | na | -0.28 | 6.04 | 6.87 | 5.30 | Platelet derived growth factor |
| 7 | 18951233 | CNOT9 | intron 3 | 0 | na | -0.34 | 4.86 | 5.65 | 7.81 | Signal transduction as well as retinoic acid-regulated cell differentiation and development |
| 4 | 61598757 | LOC102037327 | Regulatory 2 | 34311 | na | -0.41 | 4.77 | 5.55 | 9.48 | C-C motif chemokine 3-like |
| 13 | 1369943 | ARHGAP26 | intron 1 | 0 | na | 0.34 | 3.56 | 4.29 | 7.51 | Triggers cell surface receptors to begin signaling cascades that regulate the organization of the actin-cytoskeleton |
| 2 | 30058845 | HEPACAM2 | Regulatory 2 | 0 | 0.07 | na | na | na | na | Immunoglobulin superfamily regulating mitosis. Knockdown of this gene results in prometaphase arrest, abnormal nuclear morphology and apoptosis |

**Table 3.** Correlation of top candidate genes (allele frequencies) identified in FASTmrMLM and the *bslmm* model with Longitude, Latitude, hydrogen stable isotope, and the top ranked environmental variables identified in the gradient forest analyses (BIO01, BIO13, BIO15, NDVIstd).

| Chrom | Marker position (bp) | GeoFor_1.0 version 102 annotation | Region | Distance (bs) | Lon | Lat | $\delta^2\text{H}$ | BIO 01 | BIO 13 | BIO 15 | NDVI (sd) |
| --- | --- | --- | --- | --- | --- | --- | --- | --- | --- | --- | --- |
| 1A | 50451000 | PDGF-B | regulatory 1 | 2502 | ns | ns | **-0.6** | ns | ns | ns | ns |
| 4 | 61598757 | LOC102037327 | regulatory 2 | 34311 | ns | ns | ns | ns | ns | ns | ns |
| 4A | 8965887 | LOC102037655 | intron 2 | 0 | ns | ns | **-0.7** | ns | ns | ns | ns |
| 7 | 15411182 | GLI2 | intron 12 | 0 | ns | **-0.6** | **0.6** | **0.7** | ns | ns | ns |
| 7 | 18951233 | CNOT9 | intron 3 | 0 | ns | **0.6** | ns | **-0.8** | ns | ns | ns |
| 10 | 3306352 | CSPG4 | exon 7 | 0 | ns | ns | ns | **0.6** | ns | **0.7** | ns |
| 13 | 1369943 | ARHGAP26 | intron 1 | 0 | ns | ns | **0.6** | ns | ns | ns | **0.6** |

### Environmental variables associated with molting

While previous work used gradient forest to rank which environmental variables are most important to describing patterns of genetic variation across space^65,66^, here we use gradient forest to identify environmental variables associated specifically with loci underlying molting. This novel use of gradient forest allows us to focus on putative environmental drivers of local adaptation in a specific phenotype known to have important fitness effects. Overall, our GEA analysis found a strong relationship between environmental variables and genomic variation underlying molting phenotypes (i.e., the 412 variants identified in the *bslmm* analysis that had non-zero effects). In particular, bioclimatic variables did best at explaining genomic variation, with nine of the 10 most important predictors in our model representing either temperature or precipitation measurements (Fig S1-A). The top four explanatory variables in our model were BIO-15, BIO-01, NDVIstd, and BIO-13, which are precipitation seasonality, annual mean temperature, standard deviation of normalized difference vegetation index, and precipitation of the wettest month, respectively (Fig S1-B). The vegetation index, related to seasonal variance in plant green biomass accumulation, can be used to monitor the productivity cycle by characterizing the response change in patterns of vegetation cover. Spatial visualization of these variables indicated that the climate transitions to more precipitation during the wettest month with greater seasonal variation in precipitation and less variation in productivity moving from west to east along the sampling transect (Fig. S2). In addition, the central-southeastern populations were characterized by high annual mean temperatures and low variance in precipitation across the season (Fig S1-C). Further, randomization permutations demonstrated that these results were significantly different from random (p < 0.05, Fig S1-D).

Targeted genotyping using Fluidigm assays of the top-ranking gene, *GLI2* (Fig 3-A), associated with molting phenotypes in an additional 261 birds at 25 locations (12 new and 13 that overlap with the RAD-seq dataset) independently validated the association of *GLI2* and annual mean temperature (FDR-corrected P < 0.05; Fig 3-B,C). The highest allele frequencies at this SNP occurred in the southern and eastern most portion of the Painted Bunting range (Texas and the eastern cluster, Fig 3-B), areas of higher temperature, where migrants are thought to molt on the breeding ground, prior to migration to wintering grounds. The greatest variation in *GLI2*, is in the hypothesized transition zone of molting phenotypes (central Oklahoma and Arkansas).

## DISCUSSION

The timing and location of molting behavior has important fitness consequences for migratory animals. In particular, migratory birds must optimize access to food resources to survive high energetic costs associated with molting^67,68^. In addition, because feather molting leaves birds without a protective layer to buffer them from temperature extremes, there are likely strong selective pressures on choosing molting locations with minimal temperature fluctuations^69^. Here, we integrate isotopic and genome-wide genetic analyses to assess factors regulating the maintenance of variation in molting strategies in the iconic migratory songbird, the Painted Bunting. We found that the transition from molt-migrants in the west to migrants that molt on their breeding grounds further east is not coincident with a strong barrier to gene flow, running counter to hypotheses about the role of differences in the timing and location of molting in the early stages of divergence^21,70,71^. However, GWAS analysis identified several candidate genes associated with distinct molting phenotypes that are also known to be involved in feather morphogenesis and feather structure, providing the first window into the potential genetic basis of this key fitness-linked trait. Further, GEA analysis found that allele frequencies in loci linked to molt-migratory behavior are associated with environmental variables linked to precipitation, seasonality and aridity, in keeping with the push-pull hypothesis^36,72,73^. Overall, the results support the idea that locally adapted molting phenotypes have evolved in migratory birds as a means of facilitating life in seasonal environments.

A key question in evolutionary ecology is to determine whether a particular phenotypic trait leads to population genetic differentiation and ultimately speciation. Some authors have proposed that differences in the timing of molt between populations could lead to reduced gene flow across migratory divides if hybrids with intermediate molting behavior are less fit^74–76^, but empirical support for this idea has been limited^22^. Here, we investigated whether variation in hydrogen stable isotope values in Painted Bunting feathers is associated with reduced gene flow across a transect between populations with differences in the molting behavior. To address this question, we used an innovative approach that combined genomics and stable isotope analysis to reveal that the variation of the molt-migratory phenotype is not coincident with a barrier to gene flow in the Painted Bunting. Importantly, our genome-wide approach was generally in agreement with previous genetic studies, identifying four distinct genetic clusters across its breeding range, including an eastern, central, southwestern, and central southeastern population^77,78^ (Fig 1). By contrast, stable isotope analysis was in keeping with field observations and geolocator studies suggesting a shift in molting phenotypes within the central genetic group, from molt-migrants in the west to individuals in the east that molt on the breeding grounds prior to migration.

Specifically, individuals breeding in the central part of the western range exhibited larger variation in hydrogen stable isotope differential index (δ²H_diff_), suggesting that western birds experience a broader range of environmental conditions, possibly linked to moving to molting locations after breeding (e.g., molt-migrants). By contrast, eastern birds and Texas birds at the very edge of the western breeding distribution exhibited smaller δ²H_diff_ values, consistent with individuals molting in the same environments where they breed. While it is perhaps counterintuitive that molt-migrants of central part of the western range leave the breeding grounds to molt in more water-stressed regions, such as northwestern Mexico as previously demostrated^23,24,34^, it is in keeping with the idea that they are at these sites to take advantage of the monsoon period when productivity at its peak^73,79^. Thus, while overall we did not detect a barrier to gene flow coincident with a transition in molting strategies, the ability to identify molting phenotypes using stable isotope analysis opens opportunities for understanding the genetic and ecological factors underlying this important fitness-linked trait^80^.

Little is known about the genetic basis of molting behavior despite its potentially important consequences to fitness in migratory birds^81,82^. While previous genetic research has identified candidate genes linked to the speed of tail feather molt in the long-distance migratory Willow Warbler^83^, no study to date has investigated the genetic basis of molt-migratory behavior in natural populations. Here we use GWAS to identify significant associations between genetic variation and the molt-migratory phenotype and find the strongest association with a SNP located within an intronic region on the *GLI2* gene. *GLI2* is a clear candidate for involvement in feather molt, as studies have shown it is a transcription regulator of the Sonic Hedgehog (Shh) signaling pathway that specifies positional information required for the formation of adult flight feathers^84^. Moreover, *GLI2* activates and is co-expressed with Follistatin^85^, a gene that is linked to the development of hair follicles in mammals and feathers in birds as demonstrated by gene knockout experiments in *Mus musculus*^86^ and ectopic induction of feather growth in *Gallus gallus*^87,88^. In addition to *GLI2*, our association analyses identified a second gene, *CSPG4*, which has a demonstrated role in feather structure^89^. Specifically, manipulative experiments designed to reduce expression of *CSPGs* resulted in significantly thinner feathers^89^. While *GLI2* and *CSPG4* are both good candidates for playing a role in molting, further work is needed to clarify the role of these genes in the molting process. In addition, whole genome resequencing may help identify additional loci which contribute to the genetic basis of what is very likely a highly polygenic trait.

Here we take a landscape genomic approach to identify the putative environmental drivers of genetic differentiation at molt-associated loci and, in keeping with the push-pull hypothesis, we find a higher proportion of molt-migrant associated genotypes in regions characterized by higher seasonality and more extreme summer temperatures. Specifically, our GEA analysis supports the idea that the top environmental predictors of molt-associated loci include seasonality, productivity and aridity variables (BIO-15, BIO-13, and BIO-1; Fig S2). In particular, arid and low productivity environments in OK and TX are highly correlated with molt-migration linked genotypes, while higher productivity, wetter regions are associated with genotypes linked to breeding ground molting. In keeping with the push-pull hypothesis, these results support the idea that the hot and dry climates in northwestern breeding areas may place selective pressure on Painted Buntings to migrate south to molt in wetter, more productive monsoon regions of Northern Mexico (pull). Given that an estimated 52% of migratory birds breeding in this region exhibit molt-migratory behavior^36,90^, future work focusing comparisons between the results described herein and other molt-migrants may help clarify the generality of these findings.

Looking more in depth at the top candidate genes identified within our GWAS analysis, we see significant correlations between allele frequency in genes linked to molting and putative ecological drivers related to precipitation and aridity (Fig 3-B,C; Fig S2). Specifically, allele frequency in *GLI2*, our top-ranking candidate gene known to be involved in feather morphogenesis, is significantly associated with annual mean temperature. High frequency in this allele generally occurs in regions where late summer temperatures are more extreme, while low frequency occurs where late summer temperatures are less extreme (Fig 3-B, C). While false positives are a common pitfall of many GEA-based analyses^91–93^, here we are able to validate the association between allele frequency in *GLI2* and late summer temperatures by genotyping an additional 261 individuals from 25 locations and find that the significant association holds true in both the original and validation datasets. While further work is needed to understand the specific role of *GLI2* in feather production in Painted Buntings, our results suggest that molt-migratory individuals may be genetically distinct from traditional migrants at this gene and that such variation is driven by differences in summer temperatures.

In addition to *GLI2*, allele frequency in our second ranking gene, *CSPG4*, a gene known to be involved in feather structure, was significantly associated with precipitation seasonality and to a lesser extent annual mean temperature. High frequency of this gene is found in regions characterized by greater extremes between wet and dry seasons, while low frequency of this gene is found in parts of the range characterized by more consistent precipitation patterns year-round (Fig S2). Considering this gene’s known role in feather structure^89^, it is possible that the environmental conditions which are pushing migrants to molt in Mexico simultaneously result in differential selection on feather structure relative to populations which molt on their breeding grounds. Unfortunately, we were unable to successfully design SNP-type assays that could be used to test if these associations, or any of the other associations with productivity and precipitation found in our top-ranking genes (Fig S2) remained significant when additional sampling locations were added to the data set. Future research including functional characterization of these genes and other novel candidate genes would help further clarify genotype-phenotype and environmental links^94^.

## CONCLUSIONS

Here we combine genome-wide genetic and isotopic analysis to identify putative genetic and environmental drivers of variation in molting strategy in the Painted Bunting. Counter to existing hypotheses on the potential role of differences in molting behavior in the fitness of hybrid offspring, we find no support for the idea that the transition in molt-migratory behavior is significantly correlated with a barrier to gene flow. Alternatively, the results of our GWAS support the hypothesis that differences between molt-migrants and traditional migrants in molting behavior is in part controlled by genes involved in feather morphogenesis and feather structure. Further, our GEA results suggest that genetic variation in these genes is driven at least in part by differences in aridity and precipitation patterns in keeping with the push-pull hypothesis. Overall, this work strongly supports the idea that molt-migratory behavior is a locally adapted trait that has evolved to help migratory birds cope with life in highly seasonal environments.

## Acknowledgments

This work was made possible by a California Energy Commission grant to K. Ruegg, a National Geographic grant to K. Ruegg (WW-202R-17), and a grant to K. Ruegg from the National Science Foundation (NSF-1942313). Postdoctoral work conducted by A. Contina was supported in part by Jeff Kelly (NSF award 1840230). We thank the DNA Technologies and Expression Analysis Cores at the UC Davis Genome Center (supported by NIH Shared Instrumentation Grant 1S10OD010786-01) for their assistance with the Next-Generation Sequencing. Computational allocations from the Extreme Science and Engineering Discovery Environment (Xsede), as well as UCLA’s Shared Hoffman2 Cluster made this work possible. We thank Thomas B. Smith, the Department of Ecology and Evolutionary Biology, and the Center for Tropical Research at the University of California, Los Angeles, for providing laboratory and sample collection support. We also thank Eli Bridge, Tyler Michels, William Oakley, Heather LePage, Elizabeth Besozzi, John Muller, Jeff Johnson and Matt Poole for their assistance with sample collection. We credit the Painted Bunting artwork used in Figure 1 to The Cornell Lab of Ornithology, Cornell University, available at https://birdsoftheworld.org.

## Supplementary Figures

**Figure S1.**
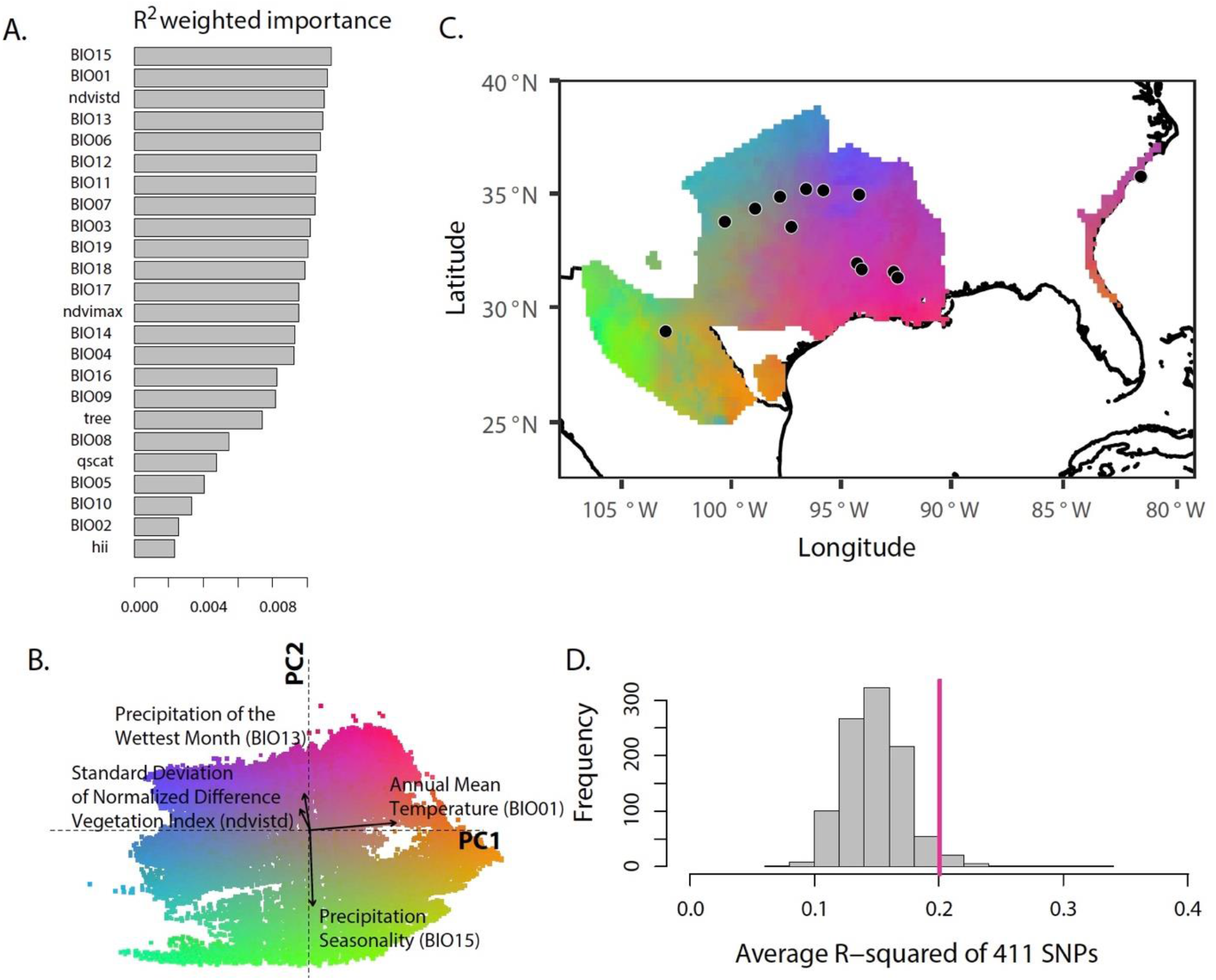
Mapping gene–environment correlations of molt-migration phenotype across the Painted Bunting breeding range. (A) Top ranked environmental variables underlying molt-migration genotypes. (B) Principal components analysis of gradient forest transformed climate variables. Colors are based upon modelled gene– environment correlations from 100 000 random points across the breeding range. Arrows show the loadings of the top-ranked uncorrelated environmental variables. (C) Gradient forest-transformed climate variables from the PCA mapped to geography support climate adaptation across the breeding range. Black dots designating approximate population locations. (D) Histogram of R^2^ of 1000 gradient forest runs of environmental variable randomizations demonstrates the variables BIO15, BIO01, BIO13 and NDVIstd are significantly correlated to the candidate molt-migration genetic variation (pink line is average R^2^ of real data).

**Figure S2.**
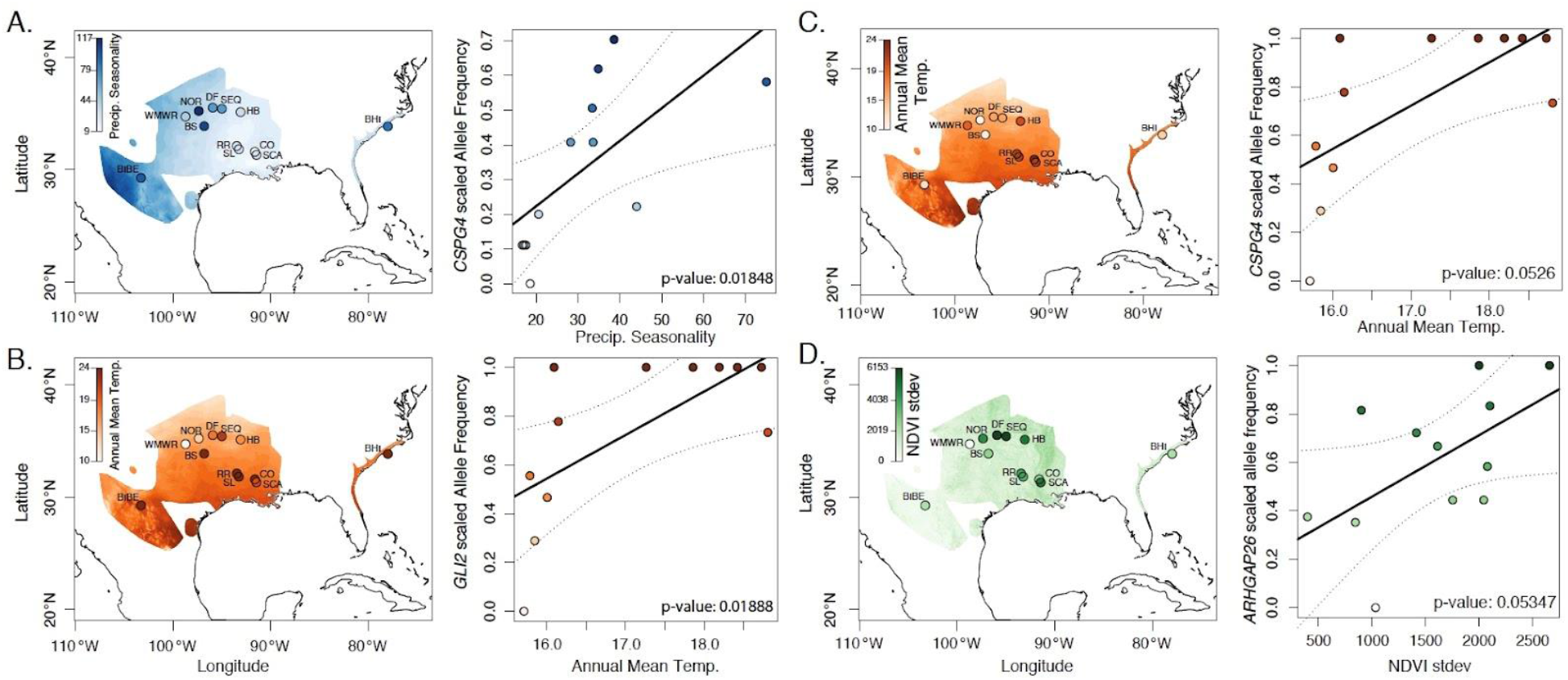
Statistical association of A) *CSPG4* allele frequency with BIO15 (Precipitation Seasonality) and B) *GLI2* allele frequency with BIO01 (Annual Mean Temperature) across 13 populations sequenced with RAD-Seq. There was moderate association of C) *CSGP4* allele frequency and BIO01 (Annual Mean Temp) and D) *ARHGAP26* with NDVIstd (variation in Productivity).

## Supplementary Tables

**Table S1**. RAD-Seq sampling locations with Latitude, Longitude, number of individuals sequenced before filtering for missingness (N_RAD_nofilter_), the number of individuals retained after filtering (N_RAD_filter_), and number of individuals for which stable isotope analysis defined molt-migration phenotype (N_Phenotyped_).

**Table S2**. List of samples used for generating the population violin plots in Fig 2 – panel A.

**Table S3**. List of samples used for generating the cluster violin plots in Fig 2 – panel C.

**Table S4**. Number of Painted Buntings successfully screened at the *GLI2* locus at each location across the species breeding range. Site ID correspond to the map code of breeding populations in Fig. 3.

## Notes

### Competing Interest Statement

The authors have declared no competing interest.

